# Ultrasonic rewarming of cryopreserved alginate encapsulated liver spheroids

**DOI:** 10.1101/2025.06.11.659069

**Authors:** Rui Xu, Tom Brookshaw, Eloy Erro, Clare Selden, Eleanor Martin

## Abstract

Rapid volumetric rewarming methods are needed to enable the effective cryopreservation and recovery of large volumes of biological cells for therapy and banking of tissues and organs. Ultrasonic rewarming is currently under development, but its effect on cells and their post-rewarming viability has not yet been established. Here, we compare ultrasonic rewarming with the gold-standard 37^°^C water bath using cryovials containing cryopreserved alginate encapsulated liver spheroids. Mean rewarming rates are used to establish the exposure time to rewarm to 5^°^C for higher power (100 W) and lower power (20 W) ultrasonic rewarming. These electrical powers correspond to free-field pressures along the central cryovial axis of 2.8 MPa and 1.3 MPa, respectively. Ultrasonic rewarming is faster than the gold-standard (120 ±5 s), taking 88 s (36% faster) and 34 s (350% faster) to rewarm to 5°C with the lower and higher powers. We measure post-rewarming liver spheroid viability and viable cell number across the 96-hour recovery period. The lower power improves viability by 1% and the higher power reduces viability by 2% on average, relative to the gold-standard. There were no significant differences in viable cell number between rewarming methods. Our findings will serve as a foundation for ultrasonic cryovial rewarming and demonstrates potential for scaling to larger volumes.

## Introduction

Ultrasound is a promising new method for the volumetric rewarming of cryopreserved media^1,2^, which is motivated by the demand for improved solutions for recovery of cryopreserved organs and large volumes of cells required for cellular therapies. Currently, the demand for organ transplantation exceeds supply^3^, partly due to the loss of viable organs which cannot be matched and used within the associated limited time frame^4^. In the long term, cryopreservation and so called banking of these organs could extend the time for optimised donor-recipient matching, but there is a still a major challenge in recovering these organs without excessive tissue damage and loss of cell viability. To overcome this challenge, improved volumetric rewarming methods are needed.

Large volume cellular therapies are being developed as curative or short term supportive treatments for a number of diseases including cancer^5–7^ and organ failure^8,9^. For example, the onset of acute liver failure is unpredictable and requires treatment within 48 hours^10^, with liver transplant often the only long-term cure. A large volume cellular therapy termed the ‘bioartificial liver device’^9^ may prolong the survivable time between acute liver failure and liver transplantation. However, the unpredictability of liver failure means that the cell volume must be made available on-demand which requires storage of the cells ready for use^9^. This may be possible with cryopreservation and effective recovery using volumetric rewarming.

The investigation of rapid volumetric rewarming methods has increased in response to initiatives to facilitate large volume cryopreservation. Several methods are under development for volumetric rewarming^11–16^. For example, a method based on magnetic nanoparticle reorientation and energy dissipation in a magnetic field termed ‘nanowarming’ has been used to rewarm 1 - 50 ml volumes of biological tissue^11,17^ including vitrified rat kidneys^14^, mouse preantral follicles^12^, porcine articular cartilage^13^. High-powered systems also show promise for nanowarming large volumes, although this work has yet to be demonstrated with biological tissues^18^. Progress toward volumetric rewarming with other methods such as dielectric rewarming^19^ is also underway, with recent work developing approaches for automatic exposure control^15^, which may reduce issues with thermal runaway. However, these methods still face challenges in the homogeneity of rewarming, due to temperature-dependent energy absorption^15^, field distribution challenges^20,21^, and nanoparticle perfusion homogeneity^11,17^. Perfusion with cryoprotectant agents that are cytotoxic at physiological temperatures (e.g., dimethyl sulfoxide) is common in cryopreservation workflows, but efforts are underway to reduce reliance on cytotoxic agents^22,23^. Perfusion with additional nanoparticles does not align with this trend and requires additional care to avoid cytotoxicity^14^. So far, demonstration of rewarming of biological samples with these techniques have been limited to volumes of less than 100 ml, below the volume of many commonly transplanted organs, and well below the litre-scale volumes needed for a bioartificial liver^9^. Further work is therefore needed to continue the development of methods for volumetric rewarming, which includes ultrasonic rewarming.

We have previously demonstrated that ultrasound can be focused through several centimetres of frozen soft tissue to generate targeted heating^2^, without the addition of any perfused exogenous agents to increase energy absorption. Ultrasonic heat deposition is proportional to ultrasound absorption, which has been shown to decrease above the solid/fluid phase transition in some biological materials^24,25^. This may enable increased uniformity of rewarming with ultrasound by preventing thermal runaway^26^ and hot-spot formation. Ultrasound has already been used to successfully rewarm cryopreserved (90% larval) nematodes from *−*80^°^C^1^ and mouse hearts from hypothermia (*−*6^°^C)^27^, but has not yet been demonstrated with cells or organoids.

Simulations have been used to plan and understand ultrasonic volumetric rewarming through low sub-zero temperatures^28^ and the phase transition^2^. However, the relationships between ultrasound pressure, rewarming rates, and post-ultrasonic rewarming cell viabilities need to be investigated to establish optimal ultrasonic rewarming regimes. There is likely to be a trade-off between ultrasound intensities that produce a significant increase in warming, reducing the risk of ice crystal growth-based damage, while avoiding ultrasound induced thermal or mechanical damage. Ultrasonic damage to normothermic cells and tissues via mechanical^29–33^ and thermal^34–36^ mechanisms have been studied in depth. Equivalent studies with cryopreserved media are needed to constrain pressure levels in future investigations.

The primary objective of this work is to determine if ultrasonic rewarming is effective in the recovery of cryopreserved cells. We evaluated ultrasonic rewarming against the current gold-standard for cryovial rewarming, the 37^°^C water bath. We selected alginate-encapsulated liver spheroids (AELS) as the cell model for this study as their recovery in large volumes is crucial for the bioartificial liver^9^. The cryopreservation protocols for AELS have been optimised^9,37–39^, but rewarming large volumes still results in sub-optimal recovery at deep thermal conduction-limited locations^40^. For example, viability at the centre of a large, thermal conduction-limited volume was 14% lower than at the edges^40^. Demonstrating equivalent AELS recovery to the gold-standard in small, non-thermal conduction-limited volumes is a crucial step toward demonstrating that ultrasound can accelerate rewarming in larger volumes while maintaining or improving viability. This study with AELS has a direct application to the realisation of the on-demand bioartificial liver, and further implications for cryovial rewarming of other cell types and for large volume rewarming.

In this work, we used a custom ultrasonic device designed for rewarming 2 ml cyrovials^41^. We completed an acoustic characterisation of the emitted acoustic pressure field and related the measured pressure amplitudes to the device driving voltage and power. Acoustic characterisation is necessary for comparison with other ultrasound devices used for ultrasonic rewarming^2,41^, and to inform the pressure amplitudes required for future large volume ultrasonic rewarming devices.

One challenge common to all rewarming methods is determining the optimal exposure time and heat deposition rate. For example, the heat deposition rate with the gold-standard 37^°^C water bath is fixed as a function of temperature, reducing the optimisation problem to one dimension, exposure time. With this method, visual determination of the disappearance of the last ice crystal is used to confirm thawing, a subjective measure that results in inter-operator variability in post-rewarming cell viability^42^. With more complex methods, variable heat deposition rates and exposure times can be implemented, giving a two-dimensional exposure parameter range that could be optimised via grid search^1^, although more efficient optimisation is desirable. Samples cryopreserved by vitrification must be rewarmed at rates exceeding the ‘critical warming rate’ to avoid ice formation during rewarming^43,44^. The critical warming rate informs the required heat deposition rate, then exposure time may be chosen to rewarm to a desired final temperature, either based on prior temperature measurement or active temperature measurement during rewarming. An alternative to freezing by vitrification is slow freezing^45,46^; this approach is used to cryopreserve AELS^9^. Slowly frozen media do not have an equivalent critical rewarming rate, but faster rewarming rates can still improve AELS recovery^40^. Previous ultrasonic rewarming work has shown improved nematode recovery at higher powers and faster rewarming rates^1^. We expect that the optimal exposure time for a given warming rate will result in a final temperature above the freezing point but below the temperature range associated with cryoprotectant agent toxicity.

In this work, we measure the exposure times required to rewarm AELS cryovials from low temperature storage to 5^°^C for a range of device powers. We then compare low-powered (20 W electrical power) and high-powered (100 W) ultrasonic rewarming with our measured exposure times, against gold-standard water bath rewarming. Our experiments show that the lower power ultrasound improves AELS viability by 0.5-1.6% while increasing rewarming rates by 36%, while the higher power ultrasound reduced viability by 2-2.3% while increasing rewarming rates by 350%. These results demonstrate promise for both small and large volume ultrasonic rewarming but indicate that careful exposure control is necessary.

## Results

### Device acoustic characterisation with free-field measurements

We modified our prototype cryovial rewarming device^2^ to improve ultrasonic transmission to the cryovial and to automate the rewarming exposures^41^. The tubular transducer radiates radially, creating a cylindrical focus at the centre of the device. Pressure measurements were performed over the volume occupied by a 2 ml cryovial when inserted into the device, as shown in Fig. 1a), the positions of the planar and line scans are also shown. Figure 1c) shows the free-field pressure (no cryovial inserted into the device) measured on a plane bisecting the transducer focus, with the transducer driven at an amplitude of 61 V. Figure 1d) depicts the free-field pressure measured along the transducer focus, with the transducer driven at voltages of 12 V to 85 V.

**Figure 1.**
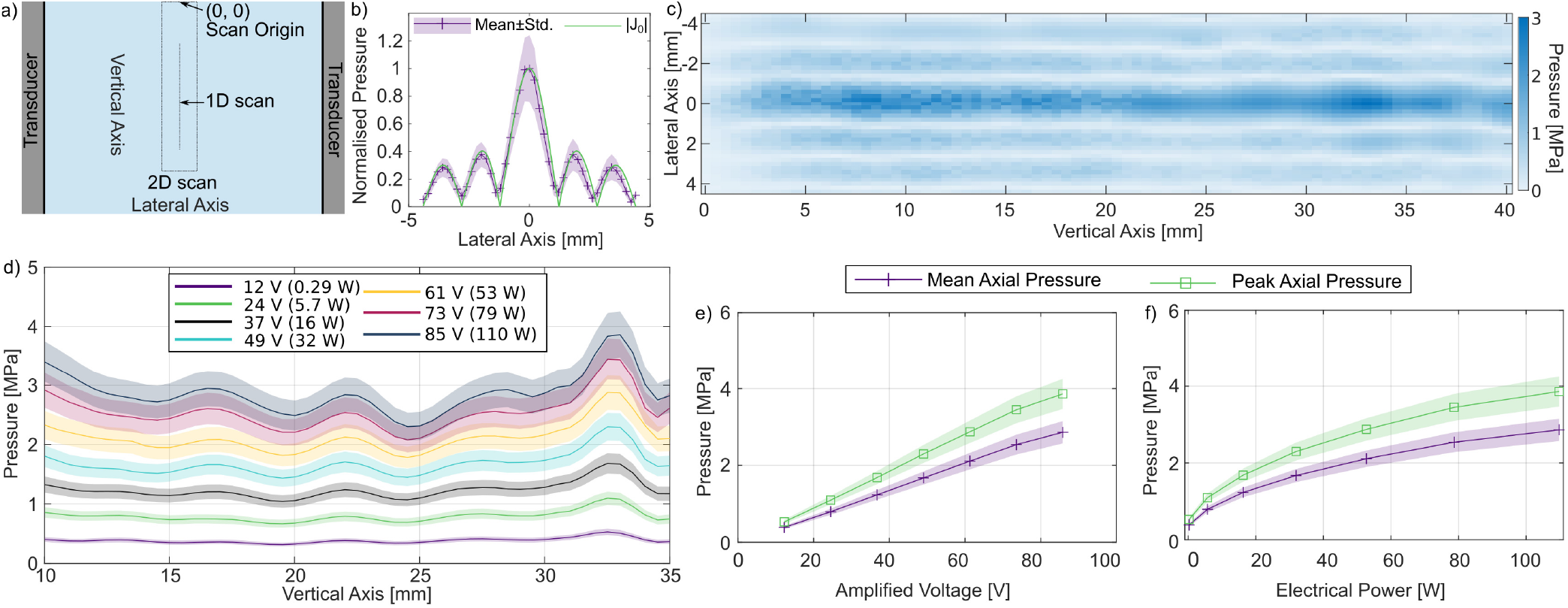
Acoustic characterisation of the device using a calibrated fibre-optic hydrophone. a) Diagram of the axes and locations of the scans displayed in b) and d). b) Mean lateral field measurement and theoretical Bessel function behaviour. The lateral field may be represented with the zero-order Bessel function *J*_0_(*kr*), where *k* is the 474 kHz wavenumber in water at 33^°^C. c) 2D measurement of the acoustic field generated in the cryovial location within the rewarming transducer, driven with a 100-cycle pulse with a voltage amplitude of 61 V, at 474 kHz. d) Measured pressure along the central cryovial axis, within the region where AELS would be positioned, for drive voltages corresponding to continuous wave electrical absorbed powers of 0.29, 5.7, 16, 32, 53, 79, and 110 W. e) Relationship between the drive voltage and mean and peak axial pressure amplitude. f) Relationship between electrical power and mean and peak axial pressure amplitude.

The device was designed to maximise the ultrasound pressure amplitude at the centre of the cryovial, where heat deposition via thermal conduction from the water bath is most limited. The radial distribution of the free-field pressure amplitude (approximating the device as an infinite tube) is given by:

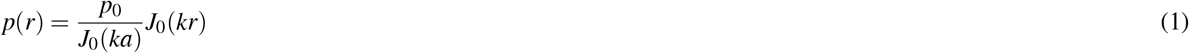

where the pressure amplitude *p* at a radial position *r* (centred within the tube) is given by the zero-order Bessel function *J*_0_, the wavenumber *k* within the cavity, and scaled by the pressure *p*_0_ and zero-order Bessel function at the tube wall *J*_0_(*ka*)^47^. Figure 1c) shows that the measured lateral pressure profiles closely match the theoretical model with *k* given by the 474 kHz driving frequency and the sound speed in water at 33^°^C^48^. The agreement between measured pressure and the theoretical model enables the calculation of the free-field spatial peak pressure amplitude from radial points within the pressure field. This approach could be used to control *in situ* pressure within the cryovial using radial pressure measurements in a similar manner to recent work in electromagnetic rewarming^15^.

The width of the central pressure lobe is directly proportional to the speed of sound in the medium. The speed of sound in frozen biological media typically exceeds that of the thawed state (e.g., by a factor of two^49^). Consequently, we expect the focal distribution in cryopreserved AELS to be broader than in water. The temperature-dependent acoustic properties of AELS and cryoprotectant solution have not yet been established, so the width of the focus in the cryopreserved AELS is not calculated here.

The planar scan in Fig. 1b) displays vertical lobes that differ from the behaviour of an infinite tubular radiator, and a higher noise level between vertical positions of 0 and 20 mm. This may be related to beat-like oscillations that originate from irregularities in the boundary condition at the top of the transducer, or due to the fibre-optic hydrophone only being partially submerged in the water bath. The vertical nodes arise from wave interactions with the top and bottom boundaries of the device, and the field is further aberrated by the introduction of the conical fibre-optic hydrophone holder into the water bath between 30 and 40 mm. Fibre-optic hydrophones can be resistant to high acoustic pressures and the tapered Fabry-Pérot polymer hydrophone is effectively omnidirectional below 1 MHz^50^. However, the acoustic radiation force^47^ on the fibre can shift the position of the sensor towards the nodes of the standing wave field, distorting the measured pressure distribution^2^. Despite these measurement artifacts, the planar pressure field broadly agrees with the theoretical distribution and demonstrates ultrasonic focusing along the central axis of the cryovial.

We measured pressure along the central axis of the transducer, between the vertical positions corresponding to the bottom of the cryovial (35 mm) and the 1.8 ml fill mark on the cryovial, at drive voltages between 12 - 85 V, corresponding to device powers of 0.28 to 110 W for continuous wave sonication. These measurements are displayed in Fig. 1d). We extracted the mean and peak axial pressures as a function of drive voltage (Fig. 1e) and absorbed electrical power (Fig. 1f). Mean and peak axial pressure increases linearly with driving voltage to 73 V, before deviating by *∼* 4% below the linear trend at the highest voltage (85 V). There is little variation in the vertical pressure amplitude profile between driving voltages when normalised by mean axial pressure; the standard deviation of the pressure amplitude was between 11% and 12% at all drive voltages. The vertical variations in pressure amplitude may result in fluctuations in rewarming rate along the vertical axis. Future work may investigate the driving frequency and the boundary surfaces as methods to reduce the vertical mode amplitudes.

The maximum peak free-field pressure that can be sustained continuously with the implemented equipment was 4.1 ± 0.5 MPa, which corresponds to a free-field spatial-peak time-averaged acoustic intensity of 560 (430 - 700) W/cm^2^, calculated using the plane wave assumption with an impedance of 1.5 MRayl, and time averaged over the whole exposure duration^51^. This peak intensity is below the 1-10 kW/cm^2^ spatial-peak time-averaged intensities used to destroy cells/tissue in high intensity focused ultrasound therapies^52^, and the peak negative pressure is below the threshold for mechanical effects in the absence of cavitation agents^53^.

### Cryopreserved AELS rewarming rate characterisation

There are several motivating factors for characterising ultrasonic rewarming rates for cryopreserved AELS. Our primary motivation for characterising the AELS rewarming rates was to obtain mean exposure times to rewarm from low-temperature storage to a desired final temperature. This knowledge will also help to develop our understanding of the physics of ultrasonic rewarming, and help to identify the possibilities and limitations of accelerated rewarming with ultrasound absorption, and will also enable comparison with other rewarming methods. Measurements of ultrasonic rewarming rates are limited^1,2^ and investigation of the factors affecting warming is still needed. Here, we investigate the effects of dead cells on rewarming rate then compare rewarming with cryopreserved AELS versus cryopreserved empty alginate beads, a possible stable AELS phantom.

In this work, we replace the final rewarming stage of an optimised and well studied cryopreservation protocol. AELS cryovials are stored in the liquid nitrogen vapour phase at *−*140^°^C, then transferred to a *−*80^°^C freezer for 40 minutes. Temperature within the cryovial is typically around *−*70^°^C at the end of this 40 minute period. The temperature distribution within the freezer is heterogeneous and cryovials stored at the front of the freezer will be exposed to warmer temperatures. The cryovials are then transferred to either the warm-water bath or the ultrasonic rewarming device (measured temperature prior to insertion: *−*65^°^C). The desired final temperature is 5^°^C, above the phase transition but below physiological temperatures to minimise cytotoxic exposure to the cryoprotectant solution^38^.

### AELS degradation slows ultrasonic rewarming rate

AELS die and degrade over successive freezing-rewarming cycles if they are not carefully recovered and re-cultured. We performed a set of experiments to determine if AELS degradation affects the ultrasonic rewarming rate via changes in the cryopreserved AELS mechanical properties. Four cryovials each containing a radially centred T-type thermocouple were filled with fresh AELS and cryoprotectant solution, then frozen slowly at −0.3^°^C/min. All experiments were completed with slowly frozen media; we found that freezing rate influences ultrasonic rewarming rates (supplementary material 1), making it crucial to match the AELS cryopreservation protocol to obtain representative rewarming rates. The cryovials were then rewarmed at an absorbed electrical power of 100 W until the thermocouple recorded a temperature of 5^°^C. We repeated the freezing/rewarming protocol three times for each cryovial. The mean rewarming curves are displayed in Fig. 2a).

**Figure 2.**
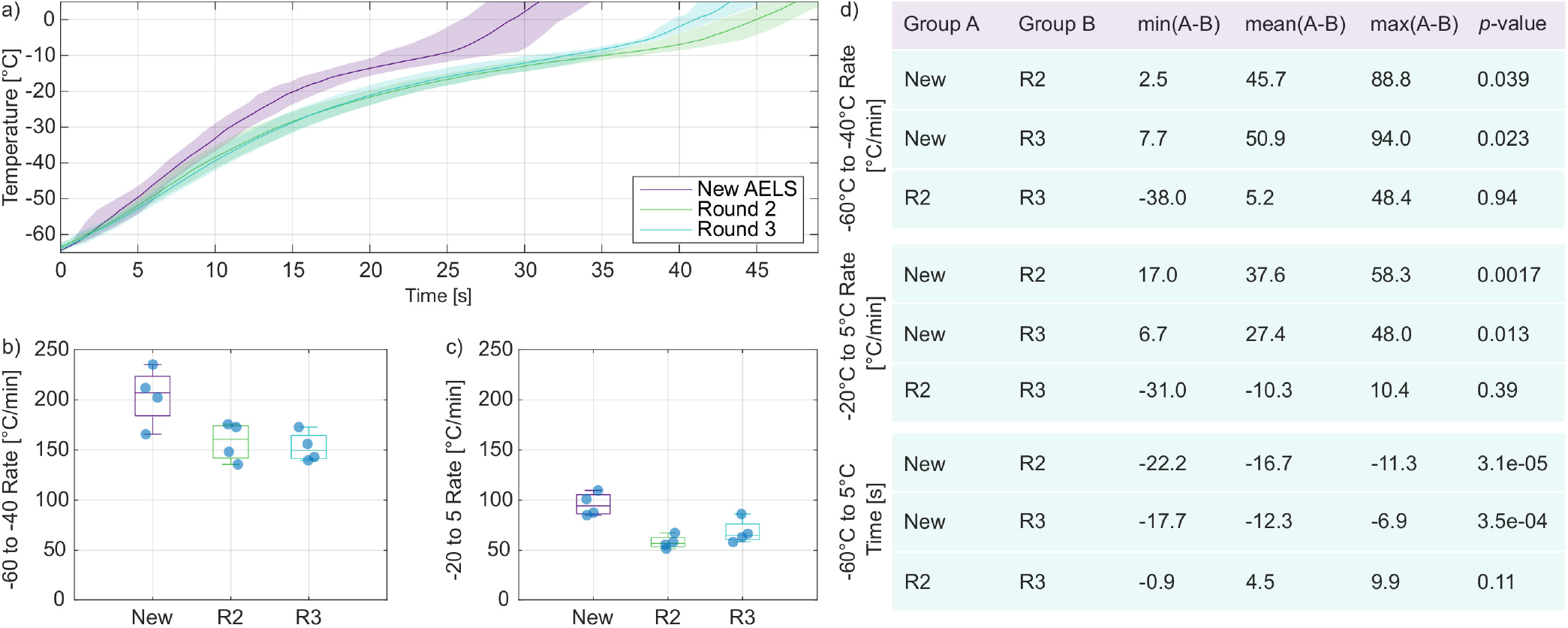
Rewarming curves for fresh versus degraded AELS. a) Mean and standard deviation in rewarming temperature for cryovials containing new alginate encapsulated liver spheroids (AELS) and old AELS which had experienced several freeze/rewarming cycles. *N* = 4 for each condition. The mean rewarming rates between *−*60^°^C and *−*40^°^C are displayed in b), phase transition rewarming rates between *−*20^°^C and 5^°^C are displayed in c). d) Results from multiple comparisons of means after one-way analysis of variance revealed differences between groups.

Mean rewarming rates were extracted from two temperature ranges, −60^°^C to −40^°^C, and −20^°^C to −5^°^C, and are displayed in Fig. 2b) and c), respectively. The lower temperature range is below the phase transition, and the rewarming rate is relatively uniform. It can be important to cross this lower temperature range quickly to avoid ice crystal growth^54^. The higher temperature range spans the phase transition between solid and fluid, encompassing changes to the acoustic and thermal properties of the medium and resulting in changes in the rewarming rate^2^. The mean rewarming rates with the cryopreserved fresh AELS were 204 ± 28^°^C/min and 96 ± 12^°^C/min for the lower and higher temperature ranges, respectively. The mean rewarming rates with the cryopreserved degraded AELS were slower. The low temperature rewarming rates for rounds 2 and 3 were 158 ± 19^°^C/min and 153 ± 15^°^C/min, respectively, and the phase transition rewarming rates for rounds 2 and 3 were 58 ± 7^°^C/min and 68 ± 12^°^C/min, respectively.

The total rewarming time from −65^°^C to 5^°^C was also extracted for each experiment. This value is the crucial exposure time for rewarming to the desired final temperature. The cryopreserved fresh AELS rewarmed to 5^°^C in 31 ± 3 s, then took 48 ± 2 s and 43 ± 4 s to reach the same temperature in rounds 2 and 3, respectively.

The degraded AELS rewarm visibly differently relative to the fresh AELS. A one-way analysis of variance confirmed that there were differences between repeats in the lower and higher temperature rewarming rates and the total rewarming time (*p* = 0.018, 0.0018, and 3*e*−5 respectively). Multiple comparison of means shows that fresh AELS rewarmed significantly faster than the ‘old’ AELS (no significant differences between repeats 2 and 3). The statistical differences between groups are summarized in Fig. 2d). This suggests that ultrasonic absorption in the cryopreserved AELS changes with repeated freeze/thaw cycles, and that repeated use of AELS in rewarming experiments will not be representative of fresh AELS. Previous work in ultrasonic microscopy has shown that ultrasonic attenuation increases in apoptotic cells at 36^°^C and 375 MHz^55,56^. Our results suggest that ultrasonic absorption in degraded cryopreserved AELS is lower than in fresh cryopreserved AELS.

### Empty beads are an acceptable AELS phantom

Characterising the relationship between device power and rewarming rate requires a large number of samples or the repeated use of a smaller number of samples over multiple freezing experiments. AELS are time-consuming and expensive to make, and are not stable over repeated experiments. However, empty alginate beads are stable across multiple freezing/rewarming cycles provided they are not heated to excessive temperatures that denature the alginate hydrogel. We investigated empty alginate beads and cryoprotectant solution as a potential AELS phantom. We performed ultrasonic rewarming at 100 W absorbed electrical power with four slowly frozen cryovials containing fresh AELS and a further four containing empty alginate beads and cryoprotectant solution. All eight cryovials were then slowly refrozen, inserted into our rewarming device and rewarmed with thermal conduction alone at 33^°^C.

The low temperature (−60^°^C to −40^°^C) rewarming rates differ significantly between the AELS and the empty bead preparation at 0 W (*p* = 0.009) but not at 100 W (*p* = 0.08). The phase transition rewarming rates (−20^°^C to 5^°^C) also differ significantly between the AELS mixture and empty bead mixture at 0 W (*p* = 0.037) but not at 100 W (*p* = 0.36) as shown in Fig. 3.

**Figure 3.**
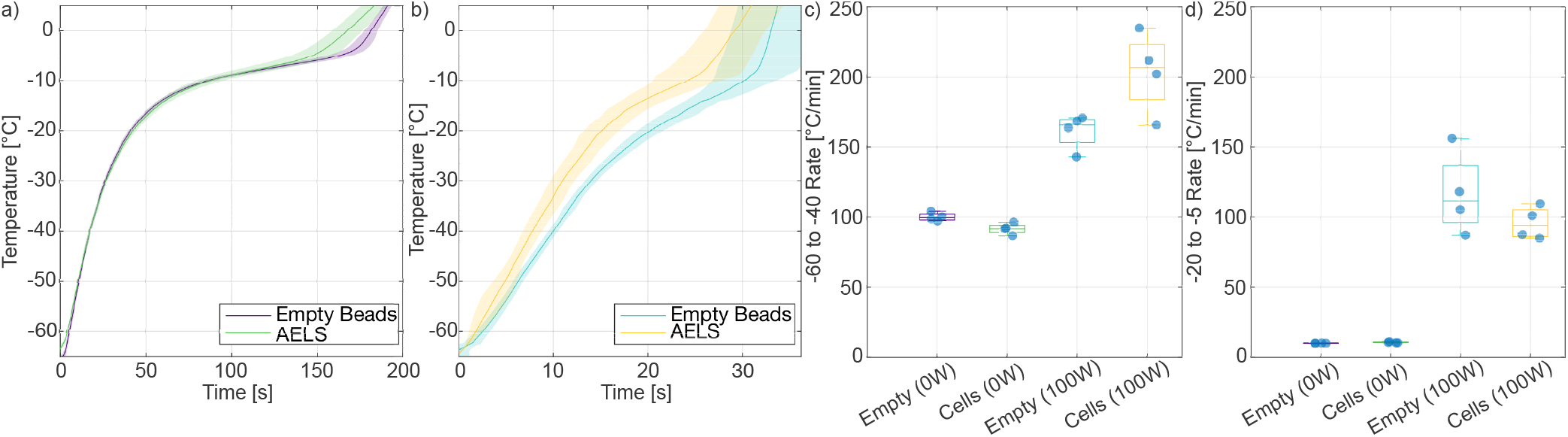
Mean ± standard deviation in rewarming curves for the a) 0 W b) 100 W ultrasonic rewarming of empty alginate beads and alginate bead encapsulated liver spheroids (AELS). *N* = 4 for each condition. The empty bead and AELS rewarming rates between −60^°^C and −40^°^C are displayed in c), while the phase transition rewarming rates between *−*20^°^C and 5^°^C are displayed in d).

The difference between the rewarming rates at low temperatures acts in opposition to the difference in phase transition rewarming rate. The *−*65^°^C to 5^°^C times for the AELS and the empty beads rewarmed at 0 W were 182 ± 7 s and 189 ± 3 s, respectively. At 100 W, the total rewarming times were 31 ± 3 s for the AELS and 33 ± 4 s for the empty beads. These total rewarming times were not statistically different between empty beads and AELS at both 0 W and 100 W (*p* = 0.17 and *p* = 0.53, respectively).

### Mean rewarming rates enable ultrasonic rewarming exposures without temperature feedback

We completed a series of experiments with *N* = 3 cryovials of empty beads to characterise the rewarming rates as a function of electrical power (0 W to 100 W in 20 W increments) and to obtain the exposure times for rewarming AELS cryovials to 5^°^C without temperature feedback control. The temperature curves generated at each power were averaged to generate mean rewarming curves and processed to obtain the temporal standard deviations at each temperature. These curves are presented in Fig. 4a). The rewarming rate of the water bath alone through the low temperature ice recrystallisation range was 100^°^C/min, equal to the commonly cited goal rewarming rate for volumetric rewarming^28,57,58^. The addition of ultrasound energy to the bath substantially increases this rewarming rate.

**Figure 4.**
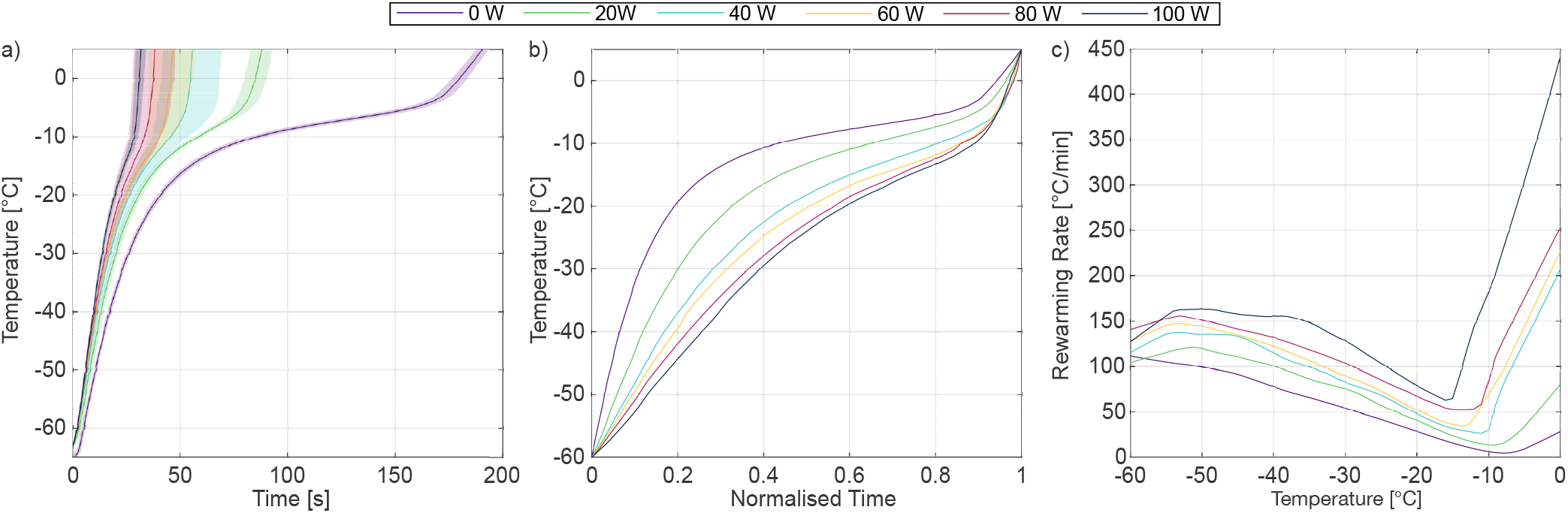
Rewarming temperature curves and rates were characterised for a range of device powers. a) Solid lines represent the mean rewarming curve and shaded areas represent the temporal standard deviation between the *N* = 3 measurements at each power. b) The mean curves from a) are normalised by their respective times to rewarm from −60^°^C to 5^°^C. c) The temperature-dependence of the rewarming rate at each power.

The tested power range spans a six-fold acceleration of the mean rewarming rate between −65^°^ and 5^°^C from 0 W (22± .2 0.5^°^C/min) to 100 W (134 ± 12^°^C/min). Figure 4a) shows that the variation between rewarming measurements in total rewarming time, a metric of reproducibility, is pyramidal. The standard deviation in total rewarming time was lowest at lower powers (±3.8 s at 0 W, ±5.9 s at 20 W) and at high powers (± 2.8 s at 100 W), and highest at intermediate powers (e.g., ±13 s at 40 W). The increase in variance between measurements at intermediate powers occurred after the phase transition, suggesting that the thermocouples were breaking out of ice at variable times during the intermediate power sonications. The low variance in the low and high powered measurements motivated the use of 20 W and 100 W sonications for the AELS viability study.

The kinetics of ultrasonic rewarming differ from water bath rewarming. Figure 4b) shows that the rewarming rate during water bath (0 W) rewarming is over 100^°^C per minute at −60^°^C, driven by the large temperature gradient between the water bath and the cryovial contents. The rate then decreases nearly linearly until the temperature reaches 8^°^C, as the temperature gradient reduces and the cryovial contents absorb the heat of fusion during the solid-fluid phase transition. The rewarming rate then begins to increase again, once the phase transition is complete and absorbed energy results in temperature rise once again.

Ultrasound accelerates both low-temperature and phase transition rewarming rates, with proportionally greater increases in the phase transition rewarming rate. The ultrasonic rewarming rates decrease through the phase transition as for water bath warming, even at high ultrasonic powers. However, this decrease is far less, with the minimum rates approximately half of the low temperature rate rather than twenty times lower as they are during water bath rewarming. The minimum rewarming rate also occurs at increasingly lower temperatures at higher absorbed electrical powers.

There are several potential mechanisms for this ultrasonic acceleration through the phase transition. Ultrasonic absorption at 474 kHz may increase with temperature through the phase transition, increasing the volume rate of heat deposition. However, attenuation measurements in frozen biological media suggest the opposite, albeit in a different material^24^. It is likely that the thermocouple viscous heating artifact^59^ becomes substantial as the cryopreserved medium becomes sufficiently fluid for the thermocouple to oscillate relative to the medium while remaining sufficiently viscous that the oscillations are damped, generating frictional heat deposition. The thermocouple viscous heating artifact can exceed temperature rise due to heat deposition from ultrasonic absorption by over an order of magnitude^41^, and may be the cause of rewarming rates that exceed 400^°^C/min at 0^°^C for 100 W. Temperature-dependent viscosity and acoustic property measurements of the cryopreserved media are needed to establish the magnitude of the viscous heating artifact on thermocouple-based measurements of phase-transition rewarming rates^41^.

Temperature measurements with metal-based thermocouples present challenges for obtaining mean rewarming rates to enable ultrasonic rewarming exposures without temperature feedback. Thermocouples measure temperature in a restricted volume and do not capture the temperature distribution across the cryovial. We measured temperature at the centre of the cryovial, where thermal conduction-based rewarming is most limited. This was in accordance with gold-standard water bath rewarming, which is based on the disappearance of the last ice crystal, which is located in the most thermal conduction limited position within the cryovial. Rewarming rates through the phase transition in the absence of a thermocouple are likely lower than those measured with an embedded thermocouple that generates viscous heating. The thermocouple viscous heating artifact is expected to be negligible below the AELS freezing point^41^. Fig. 4b) shows that at high powers, rewarming through the phase transition consists of approximately 20% of the ultrasound exposure, restricting the impact of possible artefactual heating to a relatively small portion of the exposure. If the high heating rates above the phase transition are artefactual, we would expect that an exposure time based on the measurements displayed in Fig. 4 to result in post-sonication temperatures below 5^°^C, with a notable frozen component. To assess this, we visually checked AELS cryovials devoid of thermocouples immediately after rewarming for the exposure duration required to rewarm to 5^°^C and observed either no ice or a millimetre-sized ice pellet that quickly melted. This end state matches the desired end state with the gold standard water bath rewarming method and demonstrates that our thermocouple-based temperature measurements enable ultrasonic rewarming exposures without temperature feedback.

### Ultrasonic rewarming can improve post-cryopreservation AELS viability

We completed two repeats of an experiment to assess AELS viability after rewarming with three conditions:

- 37^°^C water bath rewarming,
- ultrasonic rewarming at 20 W for 88 s,
- ultrasonic rewarming at 100 W for 34 s.

The ultrasonic exposure times were obtained from the mean rewarming rates presented in the previous section, and rounded upward to the nearest second to reflect the timing method in the exposure control algorithm. The lower power ultrasound (20 W) corresponds to a mean free-field axial pressure amplitude of 1.3 MPa and a mean free-field axial pulse-average acoustic intensity of 60 W/cm^2^, while the higher power ultrasound (100 W) corresponds to 2.8 MPa and 260 W/cm^2^. The 20 W and 100 W exposures were chosen as they evenly span the 0-2.8 MPa pressure amplitude range, and because they had smaller standard deviations in the time to rewarm to 5^°^C than the 40-80 W exposures, indicating higher repeatability. The rewarming time using the 37^°^C water bath (with agitation) was 120 ± 5 s, judged by visual confirmation of the melting of the last ice crystal.

We measured AELS viability and cell counts with fluorescence and phase microscopy immediately after rewarming (0 h), then 2 h, 24 h, 48 h, 72 h, and 96 h thereafter. The viability measurements show that all three rewarming conditions follow the same recovery trajectory, with high post-rewarming viability followed by a drop at 24 h, then an approximately linear recovery toward the initial viability. There was a difference of nearly 10% in the viability at the fourth day, between the two experiment repeats. This difference may have resulted from differences between the execution of the cryopreservation or rewarming protocol. For example, the AELS may have been frozen at differing growth phases (exponential vs. plateau, with the exponential growth phase associated with higher viability recovery). Additionally, the rewarming exposures in the second experiment repeat were completed within a shorter time (34 vs. 44 minutes to rewarm 5 cryovials for each condition). The shorter exposure time to DMSO in the second repeat may have improved the AELS recovery capacity. The cell counting protocol failed for the 20 W condition in the first repeat, but was successful for all three conditions in the second repeat. The results from this repeat demonstrate that the viable cell count for the three conditions follow the same recovery trajectory, monotonically increasing beyond the pre-freeze viable cell number of 20 million cells per millilitre by day three.

We found that the 20 W ultrasound generated a small but persistent increase of 1.6% in viability over the current gold-standard warm water bath (*p* = 0.012) in the first experiment repeat. An increase of 0.5% was observed in the second repeat which was not significant (*p* = 0.74). The 100 W ultrasound resulted in a small but persistent decrease in AELS viability (2.0-2.3%) relative to the warm water bath (*p* = 0.0014, 0.0044) across both repeats. The fluorescence image taken on day 4 for 100 W warming (Fig. 5) shows that one of the sixteen alginate beads is completely red, indicating non-viability, and supporting the measured lower viability of the 100 W ultrasound condition. Clear differences in viability between alginate beads may indicate some beads were located in a position exposed to rewarming conditions that were detrimental to AELS viability.

**Figure 5.**
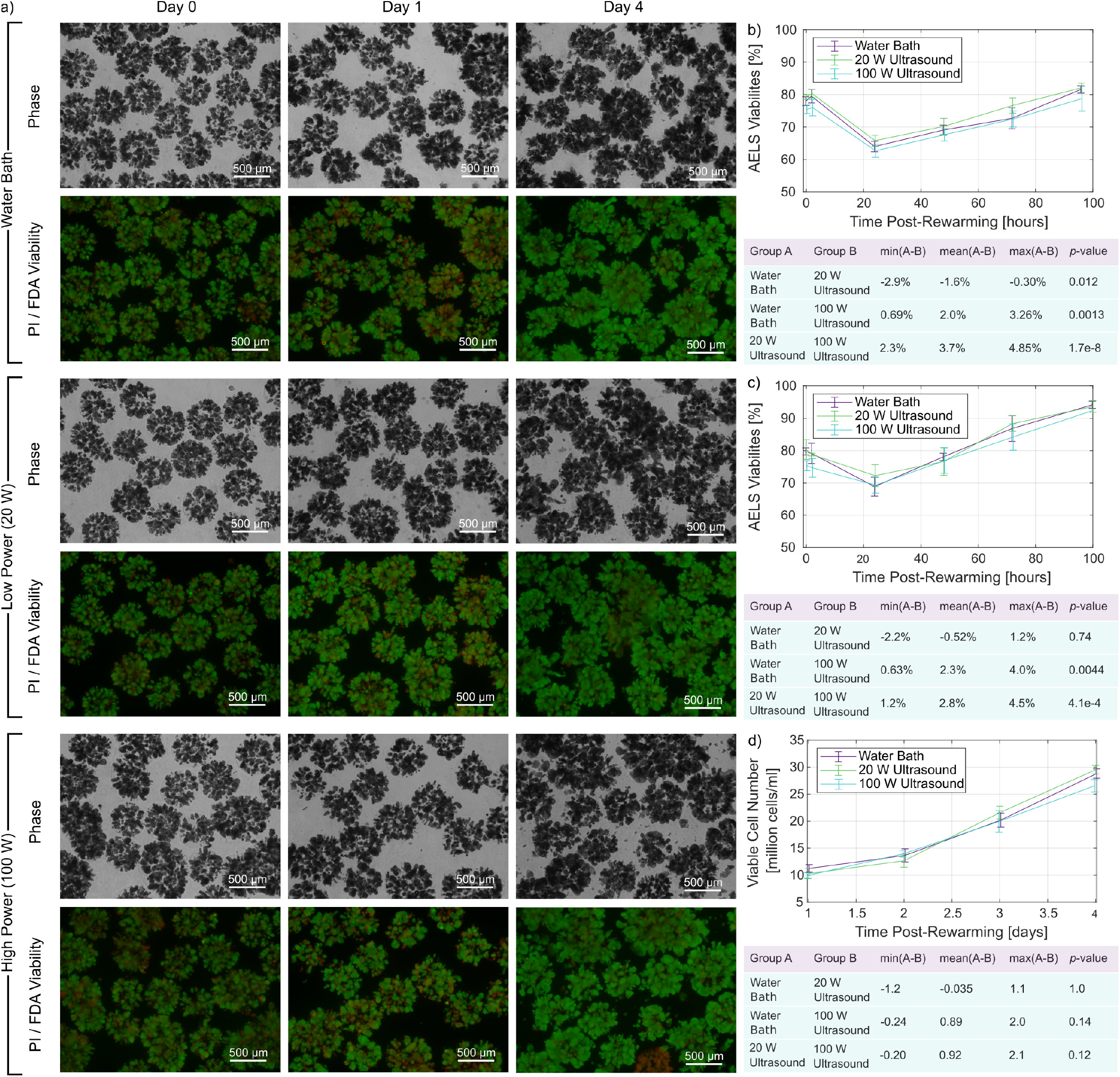
a) Alginate encapsulated liver spheroid density and viability is assessed using phase and fluorescence microscopy for: 37^°^C water bath rewarming, 100 W ultrasonic rewarming for 34 s, and 20 W ultrasonic rewarming for 88 s. *N* = 5 cryovials were rewarmed for each condition. Viability is assessed across a series of post-rewarming time-points (0, 2, 24, 48, 72, 96 hours). The viability curves and statistics for the first and second repetitions are displayed in b) and c), respectively. The viable cell number data from the second repeat are displayed in d).

The difference in viability between 20 W and 100 W ultrasound suggests there could be a threshold above which additional ultrasound intensity is detrimental to post-rewarming AELS viability. Our results differ from previous work with nematodes^1^, where their highest tested device electrical power produced the highest post-rewarming nematode viability. The device used to rewarm the nematodes was not acoustically characterised, making it impossible to compare exposure parameters, and the cryobiology likely differs between nematodes and AELS, making comparison challenging. Our viability improvements at 20 W surpass the initial small volume rewarming results with nanowarming^11^, where nanowarming matched the post-rewarming viability of the gold-standard in a 1 ml system. This demonstrates the potential of ultrasonic rewarming, and further optimisation of exposure parameters may lead to larger viability increases.

There are several potential ultrasonic damage mechanisms at high intensities and pressures including cavitation, cell membrane damage from intra- or extracellular ice crystal oscillation, and overheating or heating heterogeneity within the cryovial. Our rewarming rate characterisation and carefully selected exposure durations reduce the likelihood of overheating the cryovial contents, and the ultrasonic heat deposition reduces rewarming heterogeneity when compared to rewarming with thermal conduction alone. This makes it more likely that the lower AELS viability at the higher intensity is mechanical in origin. Cavitation is a mechanical mechanism used during freezing in the food industry, where low frequency (e.g., 40 kHz) insonation causes ice crystal nucleation via cavitation^60^. Higher frequencies and exposure control based on the real-time measurement of acoustic emissions^61^ may be useful in limiting cavitation, if cavitation is shown to reduce viability. Future work should aim to understand the origin of the slightly (2%) lower viability at the higher ultrasound intensity to further establish the regime of beneficial ultrasound intensities. Establishing this regime will be instrumental for the development of future large-volume ultrasonic rewarming devices, and to establish ultrasonic rewarming as a new method for rewarming the ubiquitous 2 ml cryovial.

## Discussion

In this work, we have presented a characterisation of our custom ultrasonic rewarming device and obtained mean ultrasonic rewarming curves that enable non-invasive exposure timing. We tested a lower and higher intensity ultrasound exposure and found that the lower intensity ultrasound slightly improved AELS viability by 1% on average and while rewarming 36% faster than the current gold-standard cryovial rewarming method. We found that the higher intensity ultrasound slightly reduced AELS viability by 2% on average while rewarming 350% faster. These findings have several implications for cryovial and large volume rewarming.

Cryovials are used across many fields of biology^62^, and even modest improvements in post-rewarming viability may be useful, particularly if ultrasonic rewarming exposures can be used to improve rewarming repeatability. Maximising viability in cryovial rewarming will likely require cell or tissue-specific exposure optimisation. We have included our device designs and control codes in the supplementary materials to facilitate the adoption and investigation of ultrasonic cryovial rewarming.

Our findings have further implications for large volume rewarming. Large slowly-frozen and thermal conduction-limited volumes have been shown to have 14% lower post-recovery viability at the centre when compared to the edge^40^, due to slow rewarming. This is a much larger drop in viability than the 2% we measured after 100 W ultrasonic rewarming. Ultrasound-accelerated rewarming rates at deep locations may improve overall viability in large volume rewarming, helping to realise large volume cell therapies.

A major concern in organ rewarming is the formation of thermomechanical stress due to heterogenous rewarming rates, large temperature gradients, and thermal expansion^21^. Thermomechanical stress can result in cracking and gross tissue damage, and is one of the primary challenges in realising organ cryopreservation and rewarming^18^. We have shown that ultrasound can substantially increase rewarming rates at depth^2^, which may be used to improve rewarming homogeneity and reduce thermomechanical stress, which may generate overall improvements in post-rewarming tissue viability. This highlights ultrasound as a promisingly versatile rewarming modality that can improve cryovial-scale to organ-scale cryopreservation recovery.

The pursuit of organ-scale ultrasonic rewarming will require the development of high-powered systems similar to those under development for nanowarming^18^. System designs may incorporate flexible and modular arrangements with half-wavelength spaced elements for electronic focusing without grating lobes^63,64^, or arrays of elements optimised for specific geometries^28,65,66^. Sophisticated arrays of hundreds to thousands of ultrasound transducer elements have been developed for the accurately targeted delivery of ultrasound energy to heat and treat a range of clinical indications^67,68^. Arrays for ultrasonic rewarming may enable the active control of the ultrasound field to obtain homogeneous rewarming rates throughout a large volume, in concert with volumetric temperature measurement or thermoacoustic simulation. Future studies should evaluate the relative merits of rapidly scanned narrow ultrasound foci for a heat deposition distribution that is homogeneous once averaged across the relevant exposure time^67,69^, or holographic methods^70^ to broaden and shape the focal distribution to deliver near-instantaneously homogeneous heat deposition distributions. Our findings suggest the latter may be preferable, to avoid the high spatial peak pressures needed to achieve an equivalent bulk rewarming rate with a scanned focus.

Our measurements span a range of ultrasound intensities and demonstrate a slight benefit to AELS viability at low intensity and slight detriment to viability at high intensity. The identified beneficial intensity and pressure is applicable to AELS, but further work is needed to determine if our findings extend to other cryopreserved media. Inspiration may be found from the field of ultrasonic imaging, which identified thermal^71^ and mechanical^72^ thresholds for damage across a range of tissue types, then developed limits for ultrasound intensities and pressures that enable safe diagnostic ultrasound imaging.

Careful exposure control is necessary to obtain a desired post-ultrasonic rewarming temperature. Our empty alginate bead phantom and thermocouple-based approach for characterising ultrasonic rewarming rates enabled the identification of exposure times for AELS recovery. This approach first required the identification of a suitable cryopreservation phantom for repeated experimentation. We found that empty alginate beads, prepared in the same manner as the AELS, were a suitable cryopreserved AELS phantom, when frozen slowly. Rewarming rate characterisations of cryopreserved media with different cell types and densities, tissue structures, cryoprotectant solutions, freezing protocols, etc., will be needed if emulating the approach taken here. We found significant differences in rewarming rates that resulted from modifications (AELS degradation, freezing rate) to the cryovial contents. It may be emphasized that these changes to the rewarming rates resulted without changing the bulk composition of the cryovial contents. Temperature measurement with thermocouples in fluid media in an ultrasound field can result in a viscous heating artifact. One modification to our approach that may reduce the viscous heating artifact is to use optical fibre-based temperature probes, which have a lower acoustic impedance that results in a smaller viscous heating artifact^59^, or to use non-invasive temperature measurement methods. Non-invasive and volumetric real-time temperature measurement may enable ultrasonic rewarming workflows to skip the rewarming rate characterisation and proceed directly to exposure control based on a desired post-rewarming temperature.

Real-time temperature measurement may simplify ultrasonic rewarming exposures, but characterising rewarming rates still provides valuable information needed for the translation of this technique to larger volumes. The combination of rewarming rate measurements with the acoustic and thermal characterisation of a cryopreserved medium will enable the validation of thermoacoustic simulations^2^. This will rely on temperature-dependent characterisations of sound speed, acoustic attenuation and absorption, density, heat capacity, and thermal conductivity, in cryopreserved media. Simulations are broadly used across therapeutic ultrasound applications to optimise application-specific array designs^66,73,74^ and complete treatment plans^74–76^. Equivalent simulation-based approaches will enable the rapid iteration and optimisation of ultrasound arrays and rewarming sonication plans for large volume ultrasonic rewarming.

Our findings are directly applicable to AELS rewarming to improve the on-demand availability of the large biomass needed for the bioartificial liver^9^, by improving post-rewarming viability of AELS rewarmed slowly via thermal conduction^40^. The findings from large AELS volume rewarming will then support future organ-scale ultrasonic rewarming.

## Methods

### Device development and acoustic characterisation

The custom ultrasonic rewarming device is based around a tubular transducer element (inner diameter: 66 mm, outer diameter: 76 mm, height: 50 mm, PI Ceramic, Lederhose, Germany)^2,41^. We added an acoustic matching layer and used deionised water as the coupling medium to improve ultrasound transmission to the cryovial, and implemented automated ultrasonic exposure control^41^. These features were added to increase rewarming rates and reproducibility and are discussed in detail in Xu *means (multcompare) was useedt al*., 2024^41^. The device is operated in a resonant, continuous wave mode during rewarming exposures, with the transducer element held above room temperature at 33^°^C. In this mode, cryovial insertion increases the absorbed electrical power during the sonication. This self-sensing characteristic was used to recognize the insertion of a cryovial and initiate a rewarming exposure for a fixed duration with a temporal accuracy of ± 0.5 s. Differences between the set and delivered electrical power are corrected using a proportional-integral-derivative algorithm to improve reproducibility^41^. We heated or cooled the device to 33^°^C prior to each rewarming exposure to initiate the sonication at the same previously optimised temperature-dependent resonant mode^41,47,48^.

We used a tapered Fabry-Pérot fibre-optic hydrophone (FOH)^50^ to acoustically characterise the device. The FOH has a 120 *µ*m fibre diameter with a 10 *µ*m active element diameter. This small sensor size and construction makes it effectively omnidirectional at low ultrasonic frequencies^50^ as well as minimising disruption to the ultrasound field.

We calibrated the FOH at 33^°^C by cross comparison with a 0.2 mm Onda HGL0200 capsule hydrophone that was previously calibrated at NPL (Teddington, UK) at 23.5^°^C. The cross-comparison was completed in a reference 474 kHz field at 33^°^C before and after characterising the rewarming transducer pressure field. We devised an experimental setup for the fast and simple transfer of the FOH between a reference ultrasonic field and the ultrasonic rewarming device. The de-ionized water in the reference tank was heated to 33^°^C using an immersion heater (Lauda, Germany), operated in the external mode using hoses connected to two radiators placed within the reference tank. Temperature within the tank was monitored with two T-type thermocouples (5SRTC-TT-TI-40–1M, Omega, Norwalk, CT), connected to a TC-08 data-logger (Pico Technologies, Corby, UK). A piston transducer (centre frequency: 500 kHz, diameter: 38.1 mm, Olympus Panametrics) was mounted at one end of the tank, and driven with a 20-cycle sinusoidal pulse at 474 kHz (200 Hz pulse repetition frequency) generated with a signal generator (Keysight 33500B, Santa Rosa, CA) and amplified by an E&I RF 75 W power amplifier (Rochester, NY) to an amplitude of 42 Volts. The signal in the far-field, 335 mm from the source, was measured with the FOH and the reference hydrophone, each positioned on laser-aligned mounts. The reference hydrophone was left to soak for 1 h before signal acquisition. The hydrophone signals were digitized with an oscilloscope (Tektronix DPO5034B, Beaverton OR) with the following parameters: 50 MS/s, a record length of 2.5k samples, and 256 averages, and saved.

Hydrophone sensitivity varies with temperature, so a sensitivity correction factor of 0.64% per ^°^C^77^ was applied to adjust the reference hydrophone sensitivity to 33^°^C from the 23.5^°^C calibration temperature. The reference hydrophone sensitivity was first used to convert the recorded voltages to pressure, then the pressure amplitude was extracted from a 10-cycle steady-state portion of the signal beginning at 231 *µ*s using extractAmpPhase from the k-Wave Matlab toolbox^78^. We then extracted the amplitudes of the FOH voltages from the same steady-state portion of the signal and used the reference pressure amplitude to obtain the FOH sensitivity before and after measurements. The uncertainty on the pressure measured with the FOH was 12%, calculated from the quadrature summation of the reference hydrophone calibration uncertainty (9% at 450 and 500 kHz) and the root-mean-square error (4%) in a linear fit to the dependence of fibre-optic hydrophone signal voltage on temperature (supplementary material 2).

We measured pressures along the central axis of the transducer and on a plane intersecting the central axis to characterise the acoustic field of the rewarming device. The transducer was driven with a signal generator (Keysight 33500B, Santa Rosa, CA) amplified by a 200 W power amplifier (E&I, Rochester, NY). Absorbed electrical power was measured with an NRT power reflection meter and NAP-Z8 sensor head (Rhode & Schwarz, Germany), used throughout to monitor power. We varied the pulse repetition frequency to maintain a transducer temperature of 33 ± 1.5^°^C, measured with a T-type thermocouple (5SRTC-TT-TI-40–1M, Omega, Norwalk, CT) connected to the TC-08 data-logger, during field scans that lasted up to 80 minutes. We used a UMS automated scanning tank (Precision Acoustics, Dorchester, UK) to position the FOH within the transducer cavity, which was filled with de-ionised water. We first aligned the hydrophone to the central peak of the pressure field. The transducer was driven with 100-cycle sinusoidal pulses to ensure cavity resonance was incorporated in the acoustic characterisation, without inducing excessive transducer heating. We completed a series of axial scans at *r* = 0 with a range of signal generator driving voltages from 50 mV to 350 mV in 50 mV intervals, amplified to 12 V to 85 V in approximately 12 V increments, to obtain the relationship between drive voltage and acoustic pressure amplitude. A pulse repetition frequency of 1 kHz was used for the lower driving voltages (50 to 200 mV) to avoid interference between pulses. The pulse repetition frequencies at the higher driving voltages (250 to 350 mV) were reduced to 600 Hz, 400 Hz, and 300 Hz respectively, to avoid excessive transducer heating. In cylindrical coordinates, we measured pressure at *r* = 0 and *z* = [10, 35] mm, with a step size *dz* of 0.5 mm, and where *z* = 0 corresponds to the top of the transducer/water level (Fig. 1a). Two-dimensional planar scans over a slice through the free-field region corresponding to the cryovial passing through the focal axis were then completed. In cylindrical coordinates, we measured pressure between *r* = [−4.4, 4.4] mm (*dr* = 0.2 mm) and between *z* = [0, 40] mm (*dz* = 0.5 mm). The dwell time at each FOH position was 1 s. The transducer was driven with an amplified sinusoidal voltage amplitude of 49 V and a pulse repetition frequency of 1 kHz. Signals from the FOH were digitized with the oscilloscope with the following parameters: 50 MS/s, a record length of 20k samples, and 256 averages, for all acoustic characterisation experiments. We found that the signal amplitudes began to stabilize after the 60th cycle and therefore extracted pressure amplitudes from the 90-100th cycles of the 100-cycle pulses to incorporate cavity resonance into the measurements.

### Preparation of cryopreserved alginate encapsulated liver spheroids (AELS)

This study used a Working Cell Bank vial containing re-derived GMP HepG2 cells (Cobra Bio, CTL2013#080P), stored in liquid nitrogen at −196^°^C. We thawed the vial in a 37^°^C water bath for 2 minutes, then used the cells to seed a monolayer triple flask (500 cm^2^, Thermo Scientific Loughborough, Leicestershire). The cells were seeded in culture media (Cytiva, S43A101351-01), and supplemented with 10% foetal calf serum (Gibco, 10500), insulin (0.27 IU/ml, Novo Nordisk, Bagsværd, Denmark), sodium selenate (Sigma, S5261), hydrocortisone (Sigma, H0888), BSA linoleic acid (Sigma, S5261), and TRH (Sigma, P1319). The cells were passaged after growing for 4-7 days, depending on cell confluency. Antibiotics were introduced to the cell culture during the second passage (45 U/ml penicillin or 45 *µ*g/ml streptomycin, Gibco 15070) and fungizone (1.1 *µ*g/ml, Sigma A2942). We changed the media every other day and increased the percentage of antibiotics by 25% each time. We expanded the cells in preparation for encapsulation after the third passage. This required seeding seven 500 cm^2^ triple flasks with 5 *×* 10^6^ cells in 100 ml cell culture media, and culturing for six days. We then harvested the cells using TrypLE Select (Gibco, 12563) and pelleted them at a relative centrifugal force of 289 (1150 rpm, Thermo Scientific Multifuge X3-R) in 50 ml Corning centrifuge tubes. We measured the viable cell number of the resuspended cells with a NC200 nucleocounter (Chemometec, DK-3450 Allerod, Denmark), then mixed the resuspended cells with alginate for encapsulation.

We prepared the alginate for cell encapsulation by dispersing and hydrating 20 g alginate as a 2% solution in 0.15M NaCl containing HEPES (Gibco, 15630, pH7.4) using an L5M-A laboratory mixer (Silverson, Ltd, Chesham, Bucks). We then encapsulated the cells at a concentration of 2 *×* 10^6^ cells/ml in 1% alginate using a Jetcutter (GeniaLab GmbH, Germany). We mixed 300 ml alginate-cell solution with a proprietary density modifier inside the Jetcutter pressure vessel, then used a cutting disc (60×100 *µ*m wires to cut the alginate-cell mixture flow into segments that dropped into a stirred polymerisation bath (0.204M CaCl_2_ solution). The pressurised delivery of the alginate-cell mixture into the encapsulator was at 400-450 mbar at 20 ml/min. We collected the encapsulated cells for 10-15 minutes, allowed polymerisation to continue for an additional five minutes, then washed the encapsulated cells in DMEM (Sigma, D6429) four times. The cells form liver spheroids once encapsulated and cultured in the 3D alginate bead environment. We produced empty alginate beads using the same protocol, without the addition of cells to the alginate solution.

We grew the AELS in twenty T175 flasks (Fisher Scientific, 10127340), each containing 155 ml proprietary media and 5 ml AELS. We used an orbital shaker (Ohaus, SHEX1619DG) placed inside an incubator (37^°^C, 5%CO_2_, RE Biotech, Galaxy R3000) to shake the flasks at 40 rpm. 80, 60, and 70% FFP media changes were completed on day 2, day 5, and day 7 respectively. Cell density was assessed at each media change with the nucleocounter. The AELS were ready for cryopreservation on day nine, with a cell density of 2 *×* 10^7^ cells/ml. We prepared a cryopreservation solution of 76% Viaspan (Belzer Cold Storage, BUWC) and 24% DMSO (Fisher, D/4120/PB08) and stored it at 4^°^C. We added two antioxidants (Trolox at 1.7 mM, Sigma, 238813, and Catalase at 500 IU/ml, Sigma, C40) to the cryoprotectant solution. We also used cholesterol (Acros Organics, 110191000) at 0.02 w/v to induce ice nucleation. We pooled the AELS from all twenty flasks in a 2 litre glass beaker in stages, removing the excess media after allowing the AELS to settle. We then placed the beaker in ice water to cool the AELS to 4^°^C before adding and mixing the cryoprotectant solution with the AELS in a ratio of 1:1 AELS:cryoprotectant solution. We again allowed the AELS to settle then removed the cryoprotectant solution until 20% excess still remained. At this stage, the AELS were ready for cryopreservation.

We loaded the AELS and cryoprotectant solution in 1.5-1.8 ml volumes into sixty 2 ml cryovials (Thermo Fisher Scientific, Roskilde, Denmark) for the cryopreservation viability assessment experiments. We then loaded the cryovials into a Kryo750 (Planer Ltd) controlled rate freezer (CRF) and froze at -0.3^°^C/min to -80^°^C. Once the freezer reached -80^°^C, we transferred the cryovials to long-term storage in vapour phase liquid nitrogen. We removed the cryovials needed for the viability experiments from vapour phase liquid nitrogen storage and placed them in a -80^°^C freezer for at least 40 minutes. The cryovials were then individually removed for the rewarming viability experiments and rewarmed sequentially.

### AELS rewarming rate experiments

We completed a series of experiments to characterise the rewarming rates achievable for AELS and to determine any effect of AELS degradation of rewarming rate and repeatability of rewarming. We prepared four 2 ml Thermo Scientific Nunc™ CryoTube™ cryovials (Thermo Fisher Scientific, Roskilde, Denmark). Each cryovial contained a T-type thermocouple with a probe diameter of 0.076 mm (Omega 5SRTC-TT-T-36-72), mounted using a 3D-printed frame to position the thermocouple tip at the centre of the cryovial, approximately 1 cm from the internal base of the cryovial. The thermocouple wire was threaded through a small hole drilled into the cryovial cap, then sealed with silicone glue (RS, 494-118). We filled the cryovials with AELS (1.8 ml) then stored them at 4^°^C in ice water. They were put into a controlled rate freezer (Asymptote, UK) once it had cooled to 5^°^C, then cooled at *−*0.3^°^C per minute to *−*70^°^C, using a heat-transfer cryovial plate. During this procedure, the cryovials supercool, resulting in a brief increase in temperature when ice begins to nucleate around *−*15^°^C.

We performed three freezing/rewarming cycles with each of the cryovials to evaluate the effect of AELS degradation on ultrasonic rewarming rate. The AELS were rewarmed with an absorbed power of 100 W, chosen to maximise the effect of acoustic absorption on the cryovial rewarming rate. This was just below the maximum stable absorbed power achievable with the power amplifier. Cryovial temperature and exposure time was recorded at a rate of approximately 2 Hz^41^. The cryovials were stored in an ice bath at approximately 5^°^C after ultrasonic rewarming, until the controlled rate freezer temperature returned to 5^°^C. The ice bath was intended to reduce higher-temperature exposure to DMSO, which can have a cytotoxic effect^38^. It does not aid recovery from cryopreservation, so the AELS were still expected to degrade across the freezing/rewarming cycles. Rewarming rates from a) *−*60^°^C to *−*40^°^C, b) *−*20^°^C to 5^°^C, and c) the rewarming time from *−*60^°^C to 5^°^C were extracted and analysed to establish the effect of AELS degradation on the low temperature rewarming rate, phase transition rewarming rate, and total rewarming time. We used one-way analysis of variance (`anova1`) to determine if there were significant differences resulting from AELS degradation between the three groups (freshly frozen, refreeze 1, refreeze 2). When the anova1 showed significant differences between groups, multiple comparisons of means (`multcompare`) was used to differentiate the group results. All statistical analysis was completed in Matlab.

### Empty bead rewarming rate experiments

One freezing/rewarming experiment was performed with each of the AELS cryovials to evaluate the AELS rewarming rate with thermal conduction alone (0 W acoustic power, 33^°^C water bath). These measurements were conducted for comparison with 0 W rewarming with cryovials containing empty alginate beads (Fig. 6), to help establish the validity of empty alginate beads as an AELS phantom.

**Figure 6.**
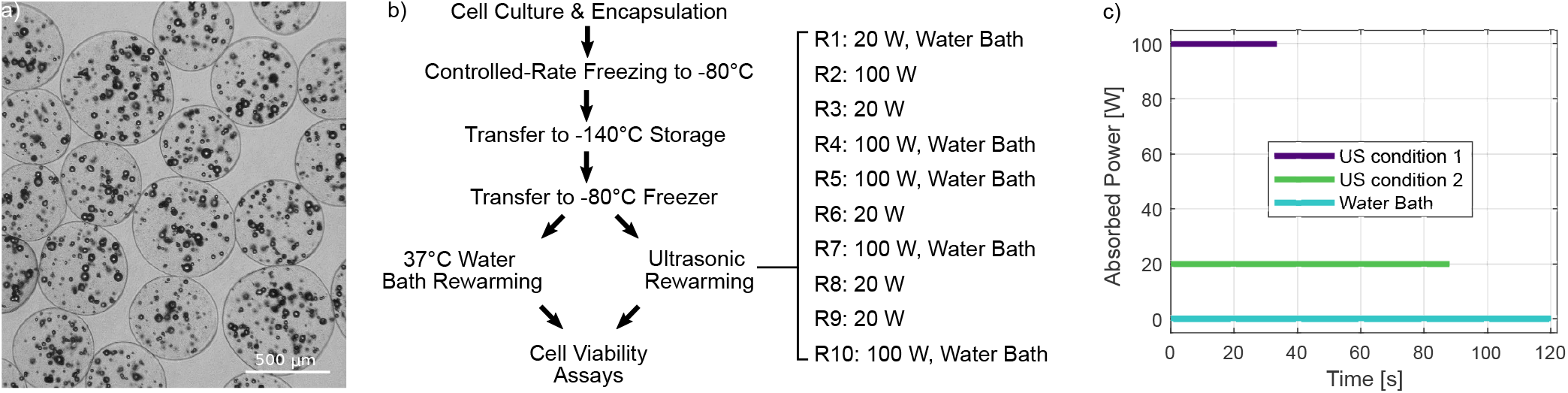
a) Empty alginate beads b) Two ultrasonic rewarming conditions (20 W ultrasound for 88 s, 100 W ultrasound for 34 s) were investigated alongside the current standard 37^°^C warm water bath, within the cryopreservation protocol. c) The ultrasonic power contributions to the three rewarming conditions.

We performed a set of experiments with *N* = 4 cryovials containing an average of 1.8 ml empty alginate beads and cryoprotectant solution to determine if empty alginate beads are an acceptable AELS phantom. Cryovials were prepared as described above and cooled slowly as previously described. The empty bead cryovials were then rewarmed with 100 W ultrasound, refrozen, and then rewarmed at 0 W (thermal conduction alone) for comparison with the AELS rewarming rates. We then used two-tailed t-tests (`ttest`) to determine if there were significant differences in the low temperature and phase transition rewarming rates, and determine if there were significant differences in the total rewarming time to 5^°^C between fresh AELS and empty alginate beads.

Empty alginate beads are stable and resistant to high acoustic pressure amplitudes, making them suited to repeated freezing/ultrasonic rewarming cycles. We used empty beads instead of AELS to investigate the range of achievable rewarming rates. We measured ultrasonic rewarming rates at 0 W (thermal conduction alone), 20 W, 40 W, 60 W, 80 W, and 100 W for *N* = 3 cryovials.

Means and standard deviations of the rewarming rates were obtained for each power, and the mean time to rewarm from −65^°^C to 5^°^C was extracted. These values were used as the rewarming exposure durations in subsequent AELS viability experiments. To improve our understanding of the temperature-dependence of ultrasonic absorption on rewarming rates, and investigate the influence of thermocouple viscous heating artifact on heating near and above the phase transition^41,59^, we also normalised the mean rewarming curves at each power by the time to rewarm from −60^°^C to 5^°^C, and calculated the ‘instantaneous’ rewarming rate at each temperature in this range for each power.

### Rewarming cell viability experiments

We tested AELS viability after rewarming with three protocols: agitation within a 37^°^C water bath until the last ice crystal melts (time: 120 ± 5 s), 20 W ultrasonic rewarming for 88 s, and 100 W ultrasonic rewarming for 34 s. Five 2 ml thermo scientific Nunc™ CryoTube™ cryovials containing AELS and cryoprotectant solution were used for each condition to obtain a sufficient AELS volume for four days of viability assessment. These experiments were completed after a set of three pilot studies, discussed in supplementary material 3.

Cryovials were stored in crushed ice (4^°^C) after rewarming to limit exposure to warmer temperatures where DMSO becomes cytotoxic^38^, before being processed together once five cryovials were rewarmed for each condition. We rewarmed the cryovials sequentially using a water bath/20 W ultrasound/100 W ultrasound scheme (Fig. 6b,c) to minimize differences between conditions in the storage time in crushed ice before beginning the AELS recovery procedure. The cumulative time spent rewarming cryovials was 44 minutes for the first experimental repeat, and 34 minutes for the second experimental repeat. The sequential protocol meant that on average, the cryovials rewarmed with the water bath and the 20 W ultrasound spent *<*1 more minute resting in crushed ice than the cryovials rewarmed with 100 W ultrasound. The small discrepancy in mean DMSO exposure between groups should not result in viability differences^38^. Transducer temperature was monitored with a T-type thermocouple (Omega 5SRTC-TT-T-36-72) and water replacement and transducer self-heating were used to ensure that the transducer temperature at the start of each sonication was 33±1^°^C.

### AELS recovery

The AELS were washed after rewarming to remove the cytotoxic cryoprotectant solution. Three tapered concentrations of glucose-modified DMEM (1M, 0.5M, plain) were used to wash, with the AELS held in a 200*µ*m mesh above a beaker to collect the wash. After washing, we transferred the AELS to sterile 50 ml centrifuge tubes and resuspended the AELS in warmed FFP media. We then divided the AELS for re-culture, growth, and viability assessment in 6-well plates. Each well was filled with 8 ml media and 250 *µ*l AELS contained in a cell strainer (Fisher Scientific, 11517532). The re-culture and growth steps were repeated as previously described for the expansion of the AELS in the orbital shaker (now oscillating at 80 rpm). Six plates were prepared for each rewarming condition, totalling eighteen 6-well plates and enabling viability assessment at six time-points. The media was changed on days 1 and 3 of re-culture by removing strainers containing AELS from the current 6-well plates and transferring the AELS to new 6-well plates with each well primed with 8 ml fresh media.

### AELS viability and density assessment

We assessed AELS viability using fluorescence microscopy and two fluorescent dyes (fluorescein diacetate, FDA, and propidium iodide, PI) to stain the cells to differentiate between viable and non-viable cells^37,38^. We transferred the AELS into a 2 ml microfuge tube (Thermo Scientific, 3434) with the volume adjusted to 250 *µ*l, washed the AELS twice with 1 ml PBS^+Ca+Mg^ (Gibco, 14040) and then resuspended in 500 *µ*l PBS^+Ca+Mg^. We then stained the AELS with 10 *µ*l FDA (1 mg/ml, Sigma F7378) to quantify the viable cells and 20 *µ*l PI (1 mg/ml, Biotium, 40016) to quantify dead cells. The staining took 90 s at room temperature. We then washed the stained AELS again and resuspended them in PBS^+Ca+Mg^ as previously described before transferring them to a microscope slide (Epredia, MIC3810) and covering them with a glass cover slip (Menzel-Glaser, 22 *×* 32mm). We used a Nikon TE200 microscope and a Nikon DS-Fi1c camera to obtain five images each for phase, FDA, and PI. The excitation for the FDA image was at 465-495 nm, the emission was at 515-555 nm, and the exposure duration was 100 ms. The excitation for the PI image was at 510-560 nm, the emission was at 590 nm, and the exposure duration was 800 ms. All images were obtained with 4x magnification and a phase objective. We quantified AELS viability from the fluorescence images using NIS Elements software (Nikon) with a custom macro^39^ that obtains the sum intensity for both sets of fluorescence images. We then determined the AELS viability as the ratio of the sum of FDA intensity to the sum of FDA + PI intensity^9^.

We assessed AELS viability at six time-points: immediately after rewarming (0 h), then at 2 h, 24 h, 48 h, 72 h, and 96 h post-rewarming. Viability time-curves are essential to cellular therapies that require a precise viable cell mass for treatment efficacy, and to assess the regeneration potential of the biomass^9^. Differences in viability between the three conditions were assessed with a two-way analysis of variance (`anova2`) after establishing the normality of the residuals using `anova` and the Shapiro-Wilk test^79^. A multiple comparisons test was then performed to determine the pairwise differences between the means of each condition, using `multcompare`. All statistical analyses were performed in Matlab.

We assessed AELS density at four time-points: 1, 2, 3, and 4 days post-rewarming. An error occurred, and we were unable to assess the AELS counts for the lower intensity ultrasound during the first experimental repeat. These data are included in supplementary material but were not analysed for differences between rewarming methods. We assessed differences in viable cell number between the three rewarming methods in the second experimental repeat using the same statistical methods implemented for AELS viability.

## Acknowledgements

This work was supported in part by a UKRI Future Leaders Fellowship (Grant No. MR/T019166/1), and in part by the Wellcome/EPSRC Centre for Interventional and Surgical Sciences (WEISS) (203145Z/16/Z). The authors wish to thank Professor Barry Fuller and Michael Brown for useful discussions on this work, Morgan Roberts for advice on assembling the acoustic matching layer, and Simon Hemsley for machining the acoustic matching layer. The authors also wish to thank Volker Wilkens and Jennifer Twiefel from the Physikalisch-Technische Bundesanstalt for extending their recent analysis of hydrophone sensitivity temperature-dependence to sub-MHz frequencies for use in this work. For the purpose of open access, the author has applied a CC BY public copyright licence to any Author Accepted Manuscript version arising from this submission.

## Author contributions

R.X., C.S., T.B., and E.M. conceived the research and designed the study. R.X. completed the acoustic characterisations and rewarming rate measurements. T.B. and E.E. prepared the AELS samples, T.B. and R.X. completed the rewarming exposures, and T.B. and E.E. completed the viability assessments. R.X. wrote the manuscript with input from all authors. All authors reviewed the final manuscript.

