## Supplementary Figures for "Ultrasonic rewarming of cryopreserved alginate encapsulated liver spheroids"

**bioRxiv preprint**

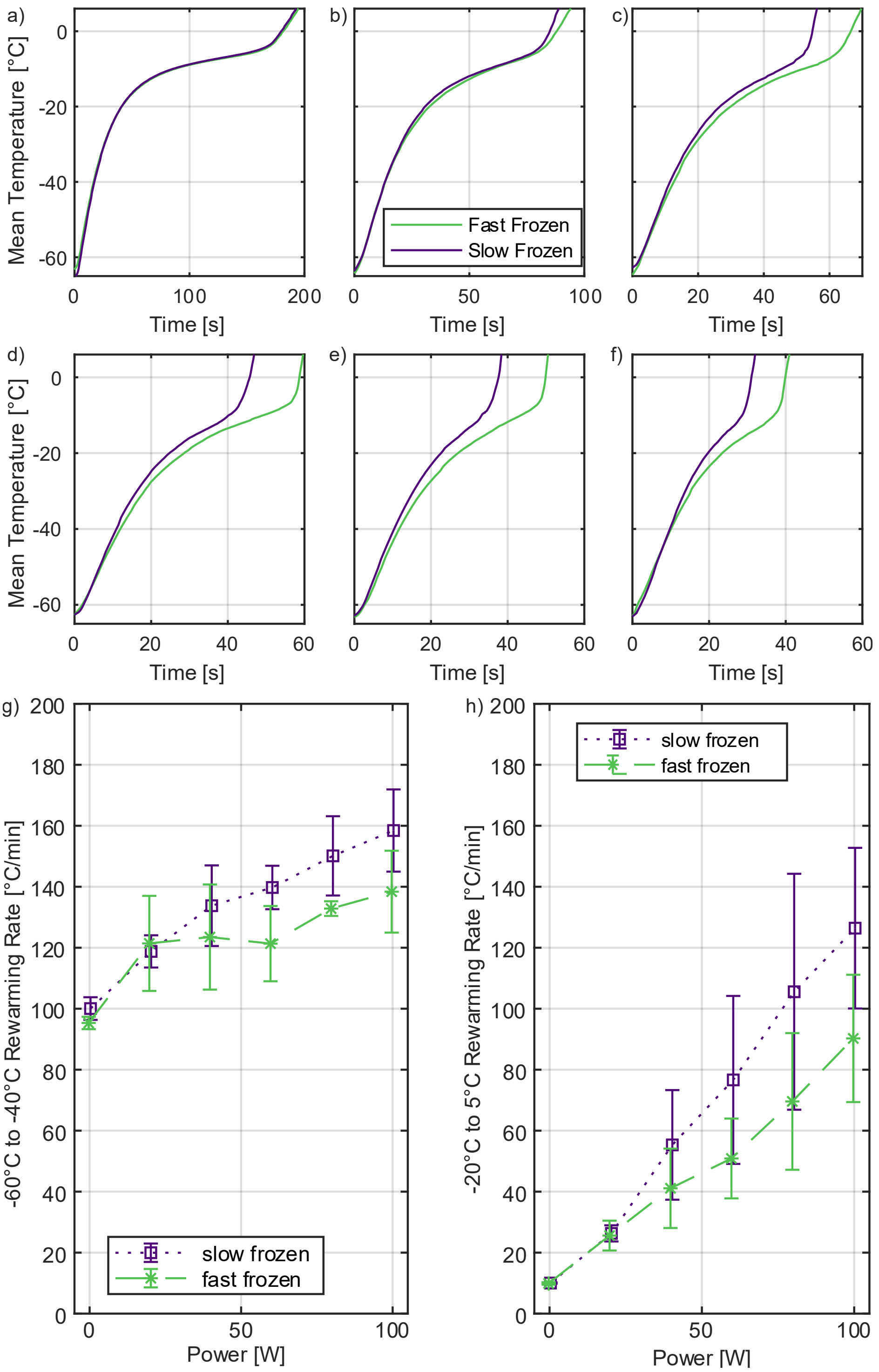


**Fig. S1. Freezing rate affects ultrasonic rewarming rate.** Mean rewarming curves for slowly frozen (−0.3°C/min) and rapidly frozen (mean of -10°C/min) empty alginate beads and cryoprotectant solution, rewarmed at (A) 0 W (thermal conduction), (B) 20 W, (C) 40 W, (D) 60 W, (E) 80 W, and (F) 100 W device power. $N$ = 3 for each condition. The mean rewarming rates diverge between slowly versus rapidly frozen media at (G) low temperatures (between −60°C and −40°C) and (H) through the phase transition (between −20°C and 5°C).

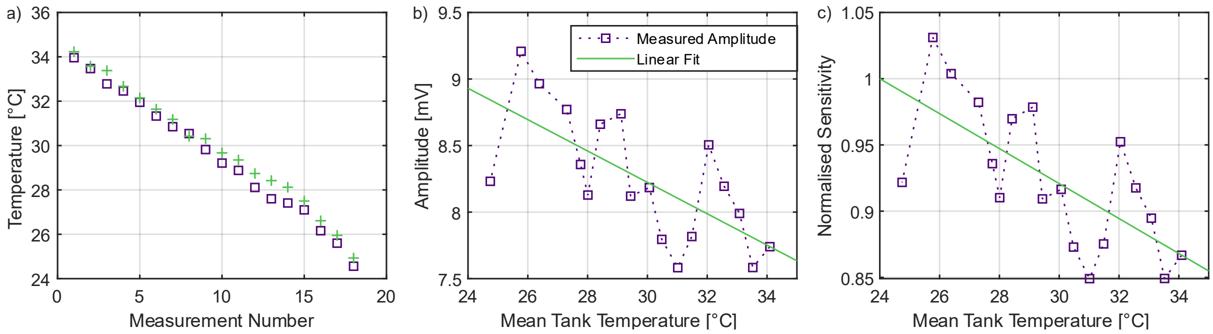


**Fig. S2. Fibre-optic hydrophone (FOH) sensitivity changes with temperature**. (A) FOH measurements were taken across a range of temperatures relevant to the rewarming experiments. (B) FOH sensitivity varies with temperature. (C) Sensitivity normalized to 24°C. To avoid issues with accounting for both FOH sensitivity temperature-dependence and transducer output temperature-dependence[80] (unknown), all characterization experiments were completed with the transducer at the operating temperature (33°C), and using a FOH calibration also obtained at 33°C using a capsule hydrophone (Onda, 0.2mm) with an expected sensitivity temperature-dependence. RMSE from the linear fit to the data in (C) was used to describe the fluctuations in FOH sensitivity.

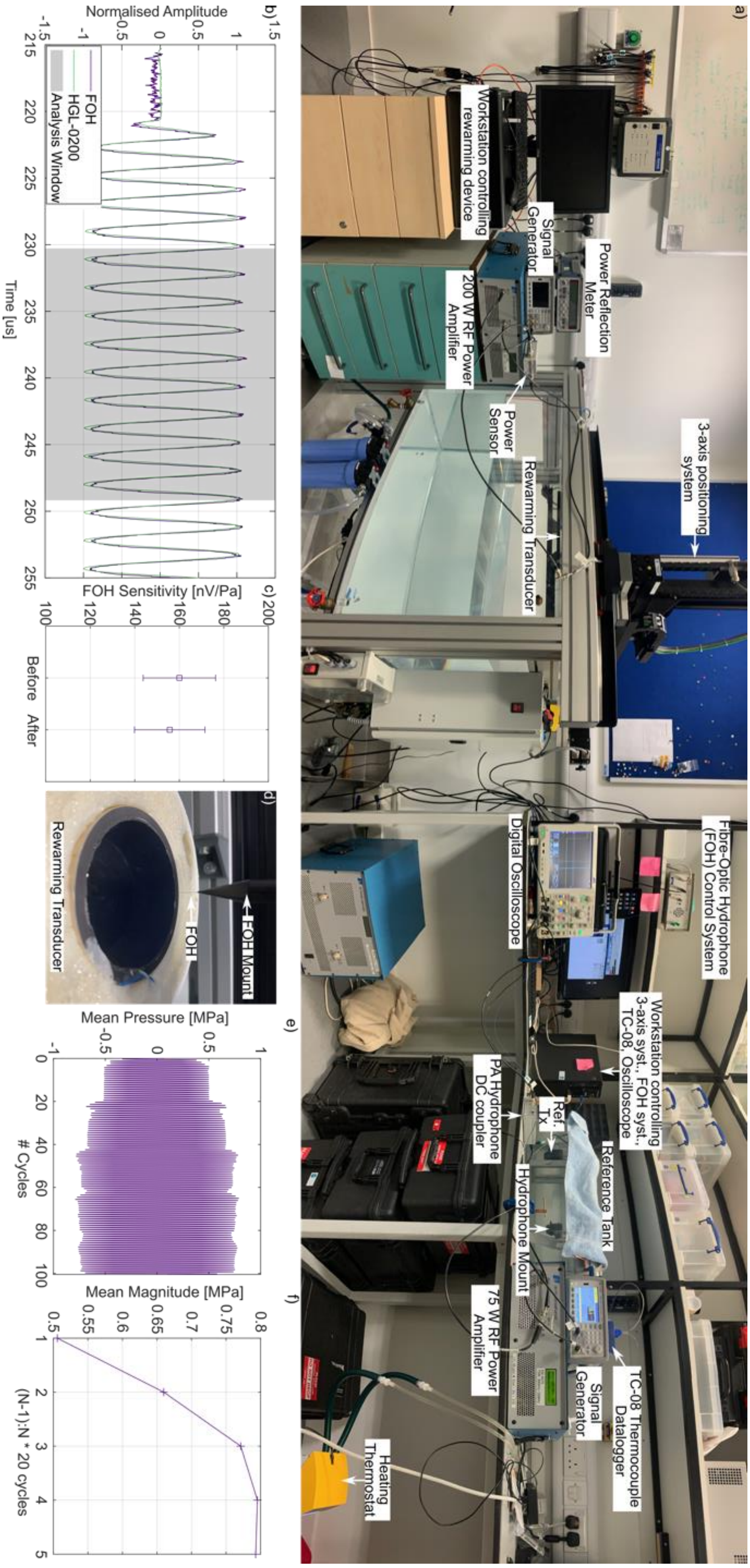


**Fig. S3. The acoustic characterization approach.** (A) Annotated photo of the experimental setup used to characterize the ultrasonic rewarming device at 33°C using a fiber-optic hydrophone (FOH). The FOH was calibrated in a 33°C reference field before and after using it to characterize the rewarming device acoustic field. (B) Normalized 474 kHz waveforms recorded in the reference tank with the FOH and with a 0.2 mm capsule hydrophone. (C) FOH sensitivity at 474 kHz before and after rewarming system characterization measurements. (D) The FOH was introduced vertically into the rewarming transducer, which contained de-ionized water. (E) The rewarming transducer was characterized with 100-cycle pulses to account for resonance within the cavity. (F) Pressure magnitudes stabilize by the third reflection (60 cycles onwards), on average, at $r = 0$ within the rewarming transducer.

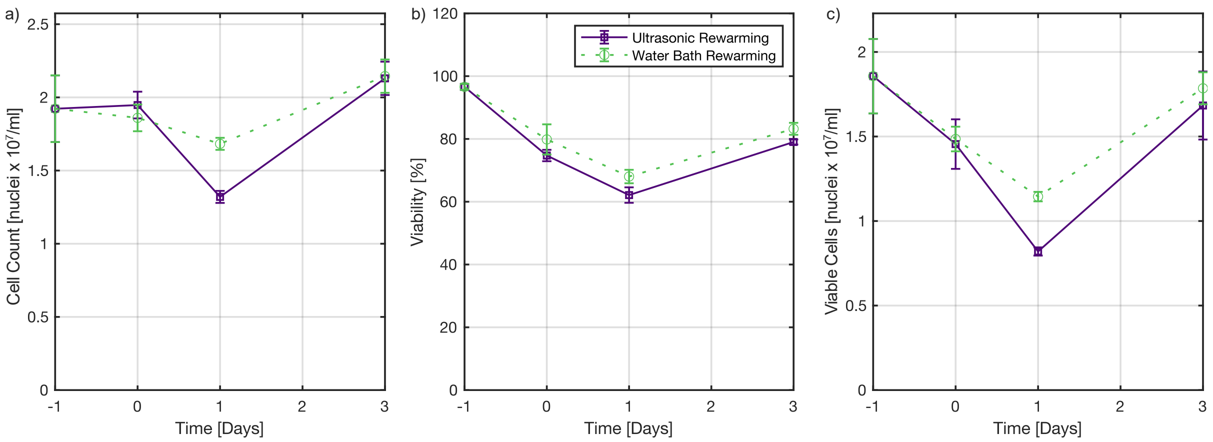


**Fig. S4. Preliminary AELS viability experiment: 80 W for 30 seconds.** (A) Cell density, (B) cell viability, and (C) viable cell density for the first round of experiments (80 W for 30 s) where the cryovials were either rewarmed via agitation in a 37°C water bath or with ultrasonic rewarming. This experiment was completed prior to the empty bead rewarming rate characterization, and the exposure duration resulted in cryovial contents that were still largely frozen (visual inspection).

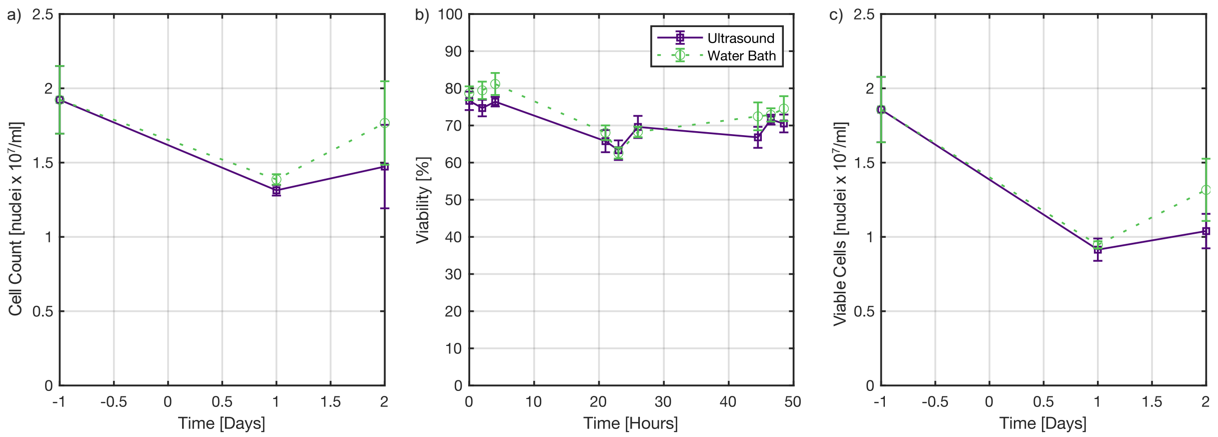


**Fig. S5. Preliminary AELS viability experiment: 100 W for 30 seconds followed by 16 W for 30 seconds.** (A) Cell density, (B) cell viability, and (C) viable cell density for the first round of experiments (100 W for 30 s followed by 16 W for 30 s) where the cryovials were either rewarmed via agitation in a 37°C water bath or with ultrasonic rewarming. This experiment was completed prior to the empty bead rewarming rate characterization, and the exposure duration resulted in cryovial contents that were fully thawed (visual inspection).

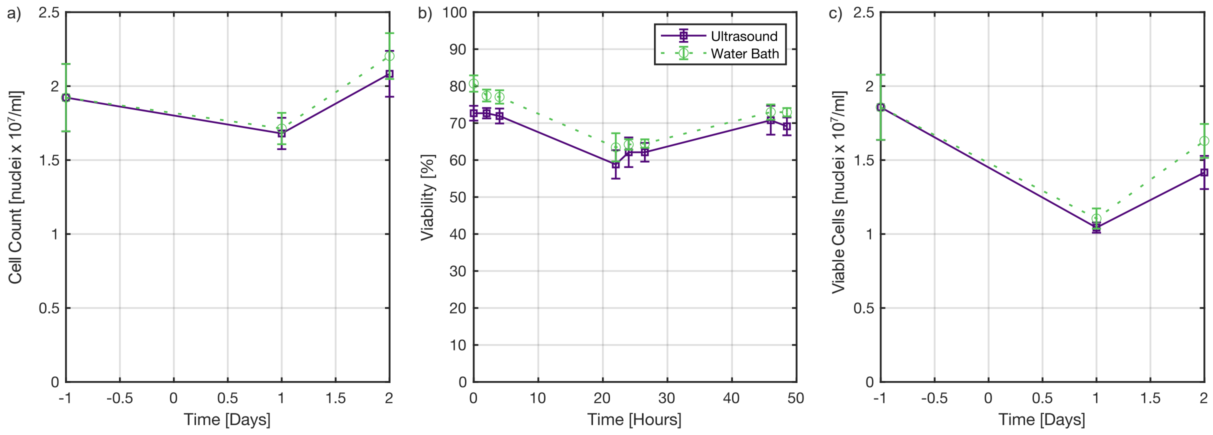


**Fig. S6. Preliminary AELS viability experiment: 90 W for 30 seconds followed by 12 W for 30 seconds.** (A) Cell density, (B) cell viability, and (C) viable cell density for the first round of experiments (90 W for 30 s followed by 12 W for 30 s) where the cryovials were either rewarmed via agitation in a 37°C water bath or with ultrasonic rewarming. This experiment was completed prior to the empty bead rewarming rate characterization, and the exposure duration resulted in cryovial contents that were mostly thawed (one 3-6 mm ice crystal visible upon visual inspection).
